# Shedding Light on functional Near Infrared Spectroscopy and Open Science Practices

**DOI:** 10.1101/2022.05.13.491838

**Authors:** Caroline M. Kelsey, Jebediah Taylor, Laura Pirazzoli, Renata Di Lorenzo, Eileen F. Sullivan, Charles A. Nelson

## Abstract

Open science practices work to increase methodological rigor, transparency, and replicability of published findings. This review aims to reflect and commemorate what the functional Near Infrared Spectroscopy (fNIRS) community has done to promote open science practices in fNIRS research and set goals to accomplish over the next ten years.

---

The open science framework is working to promote practices that strengthen methodological rigor, increase transparency in methods and analysis, disseminate null results, and ultimately, improve the replicability of published findings. Open science practices are now widely being adopted by many fields of research from basic biology to clinical and translational research (Gilmore et al., 2017; Kaplan & Irvin, 2015; Petersen, Apfelbaum, McMurray, 2022). In this special issue of *Neurophotonics*, the field is celebrating functional Near Infrared Spectroscopy’s (fNIRS) 30^th^ anniversary. On the 30th birthday of fNIRS, it is an ideal time to reflect and celebrate what the fNIRS community has done so far to promote open science practices in fNIRS research and set goals to accomplish over the next ten years. To do this, this review will go through the research life cycle of an fNIRS research project from study inception to publication – highlighting ways in which fNIRS researchers have been implementing open science research practices and ways in which the field could be doing more. The suggestions are not meant to be condemning of others’ current research practices in any way. Rather, the hope is to spark reflection and discussion about open science practices within labs and across the fNIRS field.

## Preregistration

Preregistration is a means of creating documentation of the research plan before data collection or analysis. Preregistering studies increases the transparency about what was initially hypothesized versus what was an incidental discovery and constrains researcher degrees of freedom (Simmons, Nelson, & Simonsohn, 2011; Wagenmakers, Wetzels, Borsboom, van der Maas, & Kievit, 2012). Preregistration documentation should be submitted to a publicly available repository or registry^1^. Currently, multiple user-friendly online platforms, such as the Open Science Framework (OSF; https://osf.io/), provide simple workflows for creating timestamped preregistrations. The typical preregistration plan contains the following information: Hypotheses, target sample size and power analysis, data collection procedures and measures, planned analyses, and inference criteria (for an overview and guide to the practice see Henderson, 2022). The preregistration process has several options which allow flexibility in use. These include options to write up secondary data preregistration for projects where the data are already collected but the researchers have yet to process or analyze the data; as well as the possibility to make the preregistration public immediately or after an embargo period, allowing researchers to release the information when they are ready. This process has many benefits to fNIRS researchers including a) limiting the need (or temptation) to run multiple processing streams to clean the data (e.g., by selecting specific methods and thresholds for motion correction and artifact removal), b) reducing the need (or temptation) to conduct multiple analyses (by choosing a particular analytic approach, e.g., cluster analysis, multivariate pattern analysis, functional channel of interest, etc.), c) focusing analyses on specific regions of interest d) improving organization and communication for collaborative, multi researcher endeavors, and e) aiding in the ability to publish null results and increasing the reliability of the findings. Before submitting a preregistration, it may be beneficial to run an initial pilot study or to run parameters through a simulated dataset. This can help to inform criteria selection and avoid potential missteps, including selecting criteria that are too strict (e.g., infants must look at 100% of the trials to be included) and leave you with an insufficient sample size or realizing that the data are not well-suited for the analysis plan (e.g., non-normal distribution, not enough trials due to overly-stringent inclusion parameters). Here, it is important to emphasize that preregistration is not meant to hamper scientific progress. If during the research cycle, hypotheses change due to new evidence being published or new methodologies being developed, one is not restricted from adding new measures. Rather, they are encouraged to be transparent and explicit about what was preregistered versus what was exploratory (or added *post hoc*).

### How widely adopted is this practice in the fNIRS community?

Overall, there were significantly fewer preregistrations listed on the OSF registry written about fNIRS (see *Supplementary Table 1*) compared to other widely used neuroimaging methodologies, Electroencephalography (EEG) and functional Magnetic Resonance Imaging (fMRI). However, when accounting for the total number of Google Scholar publications (used as an approximation for users across these modalities; see *Figure 1*) there are relatively similar rates of this practice. In addition, there is an upward trend such that preregistration is becoming generally more common across all three imaging modalities.

**Figure 1.**
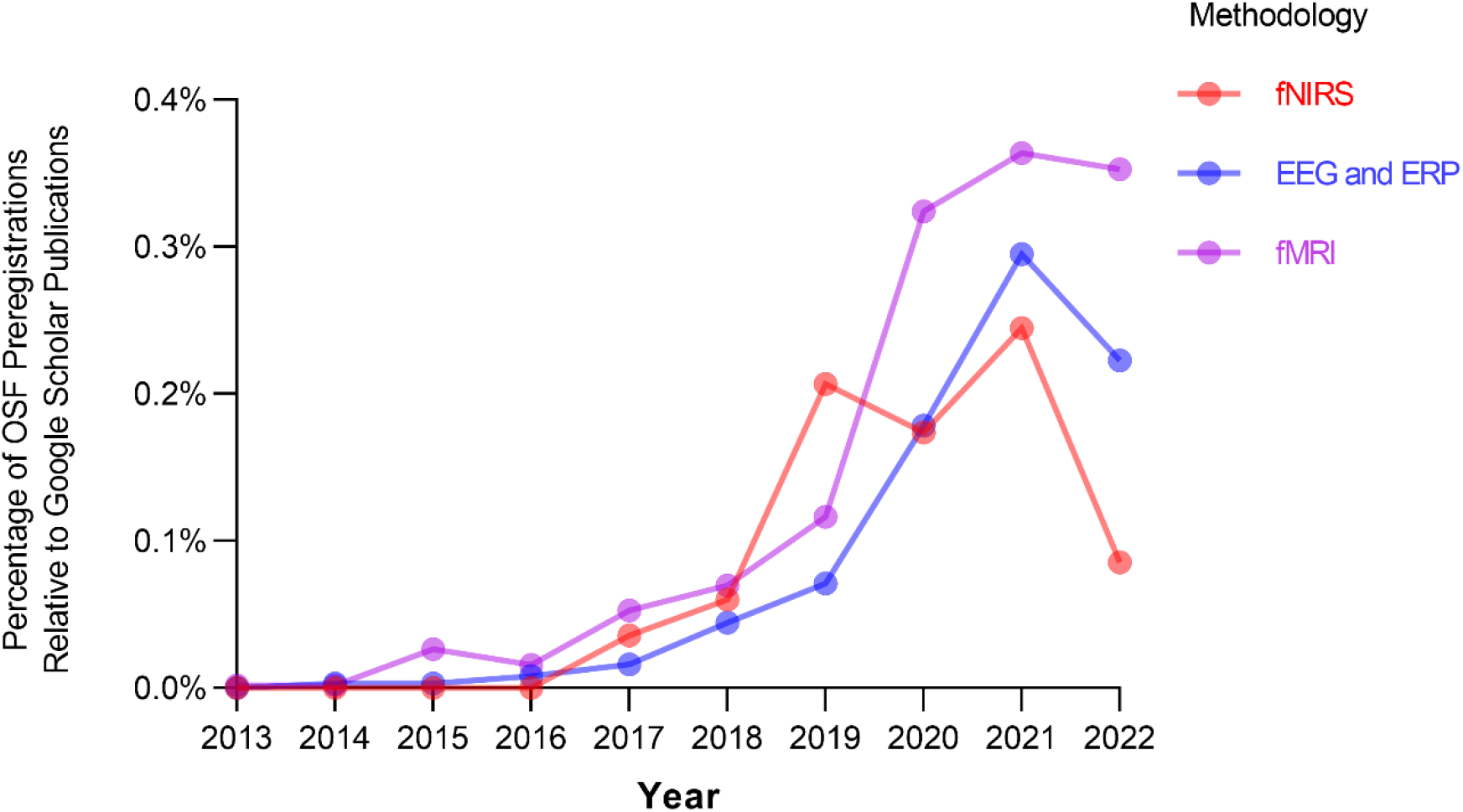
The ratio of preregistrations to publications for each imaging modality by year. *Note*. This search was conducted on 3/29/2022 using Share.osf.io (restricted to OSF Registries) and Google Scholar. This only includes preregistrations that have been made publicly available. The following search terms were used: fnirs OR “functional near infrared spectroscopy”, fmri OR “functional magnetic resonance imaging”, eeg OR erp OR electroencephalogram OR “event related potential”.

### Goals for the future

Looking ahead, the fNIRS field would benefit from creating a standardized fNIRS-specific preregistration template and an incentives program. Templates for fMRI and EEG have already been developed and can help to inform the creation of an fNIRS preregistration template (Flannery, 2020; Govaart, Paul, & Schettino, 2021). The fNIRS-specific preregistration could have a structured form field that asks about specific fNIRS hardware (e.g., the system used) and software (e.g., data acquisition software, presentation software), data processing plan steps such as behavioral coding (e.g., the minimum amount of looking time per trial needed to be included, removing periods of experimental interference), channel exclusion (e.g., channel exclusion parameters, detection of heart rate signal, etc.), motion detection correction and rejection (e.g, spline, wavelet, tpca, or pre-whitening), GLM or block average, and exclusion criteria at the trial and participant level. In addition, it would be beneficial to identify regions of interest *a priori* (e.g., description of probe layout and which channels will be used; cluster-based permutation parameters, etc.) and statistical analysis methods (e.g., functional channel of interest, multivariate pattern analysis, functional connectivity network computation, general linear mixed models). Finally, the fNIRS specific template would benefit from following the general guidelines for fNIRS research recommended by Yücel et al. (2021). We, as a field, should also reward those who are taking the extra step to promote open science practices through the creation of an incentive program (e.g., badge program). Badges are a symbolic recognition given to the researchers and they serve as a signal of values and beliefs held by the particular publisher (Blohowiak et al., 2022). Essentially, for each open science practice a paper follows (e.g., preregistration, sharing materials) they receive an additional badge (akin to a child receiving a really cool set of stickers). This easy-to-implement and no-cost badge program has shown to be effective in increasing the number of papers that endorse open science practices (Kidwell et al., 2016). Preregistration could also be incentivized through incorporation into the peer-review process by having it as part of the checklist for reviewers and something that should be considered when evaluating a manuscript.

## Registered Reports

Registered reports are a type of manuscript where the first round of peer-review occurs after the study is designed but before data are collected or analyzed (referred to as a stage 1 manuscript). Therefore, peer review at this stage focuses on the theoretical framework and quality of methods rather than the outcome of the study, which can reduce publication bias and certain questionable research practices (Chambers & Tzavella, 2022). It is at this stage that the editorial team may decide to accept (or reject) the paper, on the condition that the authors follow through with the registered methodology. After data collection and analysis, the research team resubmits the full paper (referred to as a stage 2 manuscript), and reviewers are asked to evaluate the manuscript on adherence to the protocol and whether interpretations are supported by the data. Similar to preregistrations, many journals also allow unplanned analyses to be included in the stage 2 manuscript as long as their exploratory nature is clearly stated. The registered report manuscript format has many benefits to fNIRS researchers including receiving feedback at an early stage where changes to data collection and data analysis can still be implemented. This early feedback is of considerable importance given the rapid progress in all aspects of fNIRS (from hardware to analysis tools and paradigms) and the rise of new laboratories approaching fNIRS for the first time. Hence, more experienced fNIRS experts have the opportunity to advance the field by aiding trainees and providing critical feedback on the best ways to process data and reduce researcher degrees of freedom. The registered report format may also be of interest to graduate and early career trainees as it allows one to start the writing process and receive credit on one’s CV while initial lab setup, equipment orders, IRB applications, and paradigm development are underway. Furthermore, it provides protection to researchers and promotion of replications through the promise to publish accepted projects even if other similar research emerges in the period between paper acceptance and publication of results.

### How widely adopted is this practice in the fNIRS community?

In order to assess how many journals may be open to publishing neuroimaging registered reports, the Center for Open Science (COS) registered reports database (https://www.cos.io/initiatives/registered-reports) was utilized. Here, out of the 261 journals listed on the COS website, 146 have previously published neuroimaging papers, and 44 have previously published fNIRS papers (see Supplementary Materials for more details on our methods). Note, this is not the number of journals that have published registered reports, but rather journals that may accept neuroimaging registered reports for publication given their publication history. When searching for registered report papers that have been accepted for publication, there were four research papers (see Artemenko, 2021; du Plessis et al., 2022; Guérin, Vincent, Karageorghis, & Delevoye-Turrell, 2021; Nguyen, Hoehl, Bertenthal, & Abney, 2022). The authors applaud these researchers for paving the way and hope these papers serve as a guide for those who are interested in this format.

### Goals for the future

Looking ahead, the fNIRS field would benefit from having this manuscript format more widely accepted by journals that typically publish fNIRS findings. In addition, journals should provide specific resources and checklists for reviewers to reference during each stage of review. Finally, the field may benefit from having a workshop at conferences, such as the society for fNIRS (sfNIRS) meetings, or online webinars about the registered report format to further educate about and promote this practice.

## Many labs and other multisite replicability initiatives

The many labs and other multisite initiatives are a series of collaborative replication and methodological advancements projects. The first of these initiatives was the Open Science Collaboration (OSC, 2015) where teams of researchers across the globe attempted to replicate 100 studies from the field of psychology. They found that only 36% of the replications had significant results which calls into question both the original methods and findings. This is very concerning given that current hypotheses, grant applications, and research programs stem from ‘old’ and established effects and methodologies. Since then, several neuroimaging specific replication and reproducibility studies have been conducted (Botvinik-Nezer et al., 2020; Li et al., 2021; https://www.eegmanypipelines.org/) including a few ongoing efforts for fNIRS. The major theme coming out of the neuroimaging replication efforts is that even seemingly small decisions at various points in this process can substantively alter end results (Gilmore et al., 2017; Li et al., 2021). Overall, these results emphasize the need for more replication-based multisite initiatives alongside collaborative efforts to promote robust methods.

### How widely adopted is this practice in the fNIRS community?

Here, two current initiatives being made in the fNIRS community will be highlighted. The FRESH fNIRS Reproducibility Study Hub (https://openfnirs.org/data/fresh/) is a multi-lab initiative currently being launched by Robert Luke, Meryem Ayşe Yücel, and Rickson Mesquita. This project will provide participants with two fNIRS datasets and will ask participating teams to process and analyze the data as they see fit. The overall goal is to understand the variability in fNIRS processing and analysis strategies used across the field, and the potential consequences this has on data interpretation. Another initiative, Many Babies, is working to replicate developmental findings. One of these projects, ManyBabies 3 NIRS (MB3N), is a collaborative effort led by Judit Gervain to use fNIRS to understand the mechanism by which infants learn and apply rules (https://manybabies.github.io/MB3N/).

### Goals for the future

Looking ahead, it will be exciting to see the results of the current fNIRS many labs initiatives. In addition, the hope is that these collaborative efforts will bring the fNIRS community closer together, get more fNIRS researchers involved, and create the necessary infrastructure to support future cross-site initiatives. As the field grows and there are a larger number of experts and sites with the necessary resources and equipment. Hence, fNIRS could be integrated into other large-scale studies such as Adolescent Brain Cognitive Development (ABCD; https://abcdstudy.org/) and Healthy Brain Child Development (HBCD, 2022) which have thus far focused on EEG and fMRI data collection due in part to these methodologies having a wider user base. Moreover, thanks to rapid technology developments, full head coverage systems are now available from an increasing number of companies, further aiding replication efforts. Indeed, variability in number of available optodes across systems can hinder replication of findings by imposing limitations on array design and choice of cortical regions to interrogate.

## Sharing materials

Sharing materials can take many forms including sharing stimuli, paradigm presentation scripts, preprocessing streams, and analysis code. In the 30 years of fNIRS research, paradigms, preprocessing streams, and analyses have become increasingly complicated and reliant on in-depth programming knowledge (Bowring, Maumet, & Nichols, 2019). Requiring such a knowledge base creates barriers to entry for new researchers and is even prohibitive to more experienced researchers who relied upon more traditional methods. Therefore, for the field to progress, the methods must be widely disseminated and in a user-friendly (step-by-step) manner (see Obels, Lakens, Coles, Gottfried, & Green, 2020 for tips on code sharing). Additionally, sharing materials promotes replicability and aids advanced researchers to refine or build new methods inspired by others in the fNIRS community. Sharing such tools and materials (see Mazziotti et al., 2022; Meidenbauer, Choe, Cardenas-Iniguez, Huppert, & Berman, 2021 for examples of sharing fNIRS presentation paradigms) has become easier across all fields with code sharing databases such as GitHub, and fNIRS specific code sharing databases (e.g., https://fnirs.org/resources/data-analysis/software/). In addition, our fNIRS journal, *Neurophotonics*, already promotes code sharing through its acceptance of tutorial manuscripts and also provides suggestion for code sharing on its website (see https://codeocean.com/signup/spie).

### How widely adopted is this practice in the fNIRS community?

Overall, there were substantially fewer repositories shared on GitHub about fNIRS compared to other modalities (see *Figure 2* and *Supplementary Table 2*).

**Figure 2.**
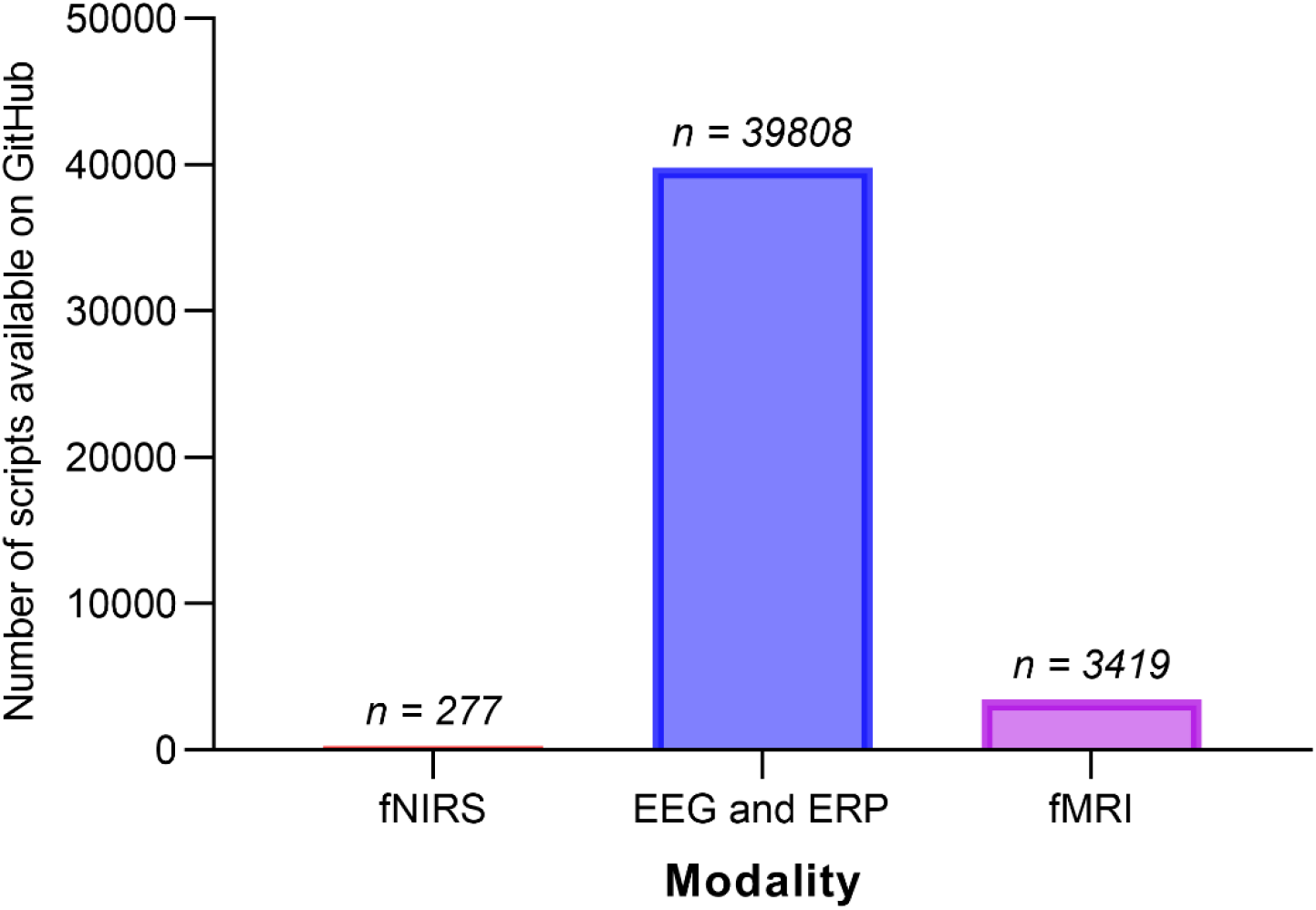
The number of codes available on GitHub for each imaging modality. *Note*. This search was conducted on 3/29/2022 using GitHub. The following search terms were used: fnirs OR “functional near infrared spectroscopy”, fmri OR “functional magnetic resonance imaging”, eeg OR erp OR electroencephalogram OR “event related potential”.

### Goals for the future

In the future, the sharing of materials can be incentivized with a similar strategy to that of preregistration, with the use of badges and having them be a criterion for reviewers to assess when making suggestions for publication. In addition, the field can work to have an agreed-upon format for code sharing. Journals may even want to go one step further by expecting code sharing but allowing for reasonable exceptions. Moreover, fNIRS researchers can continue to strive to make fNIRS research more accessible through the use of freely available software (e.g., using Psychopy for stimulus presentations, developing programs -like HOMER - that can be used on MATLAB Runtime without requiring a license, creating fNIRS processing packages for R or Python, etc.). In addition, more support for programming and materials sharing can be provided through hosting educational sessions (such as HOMER3 tutorials and sfNIRS educational events at conference meetings) and creating coding curricula for undergraduate and graduate studies. Another option for sharing protocols is through sharing videotapes of testing sessions. This may allow secondary data users insights into slight nuances of the paradigms and testing conditions that are difficult to glean from testing manuals (Gilmore & Adolph, 2017). Finally, sharing of materials can be encouraged by sponsoring calls for papers with tutorials focused on advanced fNIRS processing and methodologies that solve common issues faced by specific fields (see *Developmental Cognitive Neuroscience’s* special issue, “EEG Methods for Developmental Cognitive Neuroscientists: A Tutorial Approach” for an example).

## Sharing data

Data sharing refers to posting raw fNIRS data to publicly accessible repositories and this practice is of great importance because it facilitates secondary data analysis and meta-analytic efforts. Moreover, data sharing can help to improve the reproducibility of research by allowing others to identify mistakes and provide suggestions for improved analyses. There are a variety of online repositories that support the storing and sharing of functional neuroimaging data^2^. Because the structure and format of fNIRS data can vary across acquisition systems, processing streams, etc., the Society for fNIRS has proposed the Shared Near InfraRed File (SNIRF) as the official file format for fNIRS data (https://fnirs.org/resources/data-analysis/standards/ and https://github.com/fNIRS/snirf/tree/v1.0). SNIRF is a standard, universal, HDF5 format supported by a variety of common languages and programs (MATLAB, including HOMER3; Python, including MNE, etc.). Another effort at standardization involves the creation of a Brain Imaging Data Structure (BIDS) extension specific to fNIRS data (and utilizing the SNIRF format), which as of publication is nearing its final stages of review and release (https://github.com/bids-standard/bids-specification/pull/802). The creation of the BIDS format will be especially welcome, as platforms like OpenNeuro require data to be BIDS-compliant. The authors are extremely grateful for the various contributors to these efforts, and we echo their recommendation for adopting SNIRF as the standard format for sharing fNIRS data. Finally, across data platforms, there are debates as to if data should be shared in the raw or processed (e.g., concentration change values for each condition already computed). Therefore, there is a need for consensus in regard to data sharing practices.

### How widely adopted is this practice in the fNIRS community?

Currently, there are 11 data files available on the openfNIRS website (https://openfnirs.org/data/) and 7 available on openfnirsdb.org. This hails in comparison to the number of datasets available for fMRI (*n* = 488) and EEG (*n* = 81) on Open Neuro (https://openneuro.org/), alone (see Jiao, Li, & Fang, 2022 for an examiniation of the frequency of data sharing across fields).

### Goals for the future

In the future, formal guidelines for sharing data (e.g., file format, data structure, raw vs processed) can be created. By creating a uniform structure, researchers may be able to more easily merge datasets that are differing in acquisition systems and probe layouts. In addition, the sharing of data can be incentivized by recognizing researchers who have posted their data through a badge program and having it be a criterion for which reviewers assess a manuscript (see also this editorial on recognizing authors of public datasets, “Time to recognize authorship of open data”). These efforts will support future meta-analytic efforts and will further inform innovation and analytic tools. Moreover, these efforts can also support student or early career researchers that may not have the resources to collect their own data but have an interesting theoretical question that can be asked using someone else’s data. In addition, data sharing platforms should continue to strive to become increasingly accessible and streamlined into the everyday research cycle so there is not too large of a time or financial (e.g., requiring a full-time data manager) burden on the researcher. Importantly, platforms should be designed in such a way that they are well suited for all stages of one’s career, such that each platform is easy to use regardless of the dataset size (e.g., a single condition fNIRS study with five participants or a multi-site collaboration with thousands of participants). Finally, there are important ethical and financial considerations that can arise when sharing data. Specifically, researchers must ensure that participants are fully informed about the scope and specificity of the data that will be shared, as well as the public nature of online repositories (see Databrary, Simon et al., 2015, for templates for consent form language for sharing identifiable data with other researchers). Moreover, users of secondary data must also be held accountable for handling identifiable data with care (APA data sharing working group, 2015). Sharing of identifiable data (zip code, photos, job description, birthday, etc.) may create the potential for sampling biases such that people from historically disadvantaged groups may feel less comfortable with sharing data. Therefore, extra care and communication with the participants is needed. In addition, navigating issues of data ownership and data sharing can be difficult depending on government, institutional, and grant policies. Therefore, it is recommended to develop educational resources for researchers (e.g., templates that use easy to understand and thoughtful language for consent forms for the release of fNIRS data), host outreach programs to help educate the public on costs and benefits of data sharing, and host sessions at sfNIRS where panelists discuss ways in which researchers have been successful (or not so successful) in navigating issues related to ethics, data ownership, and data sharing.

## Preprints and open access

A preprint is a complete manuscript that you share with a public audience without having undergone peer review. Preprints allow for findings to be shared expediently and without a cost to the reader (Sarabipour et al., 2019). One potential downside of a preprint is that it contains identifying information that can bias the peer review process. A similar movement is being made by journals by making some articles available as open access. *Neurophotonics* has set an example for the fNIRS field by making all articles open access. One downside of open access publishing is that the financial burden falls on the authors of the papers. This can be prohibitive for some smaller research groups and trainees.

### How widely adopted is this practice in the fNIRS community?

To quantify how fNIRS is doing relative to other methodologies a search was performed using two popular preprint sites, *bioRxiv* and *PsyAriv*. Overall, there were significantly fewer preprints written about fNIRS (see Supplementary *Table 3*) compared to other methodologies, and this pattern held even when accounting for the total number of publications on Google Scholar (used as an approximation for users; see *Figure 3*). In addition, there was an upward trend such that preprints are becoming generally more common across all three imaging modalities.

**Figure 3.**
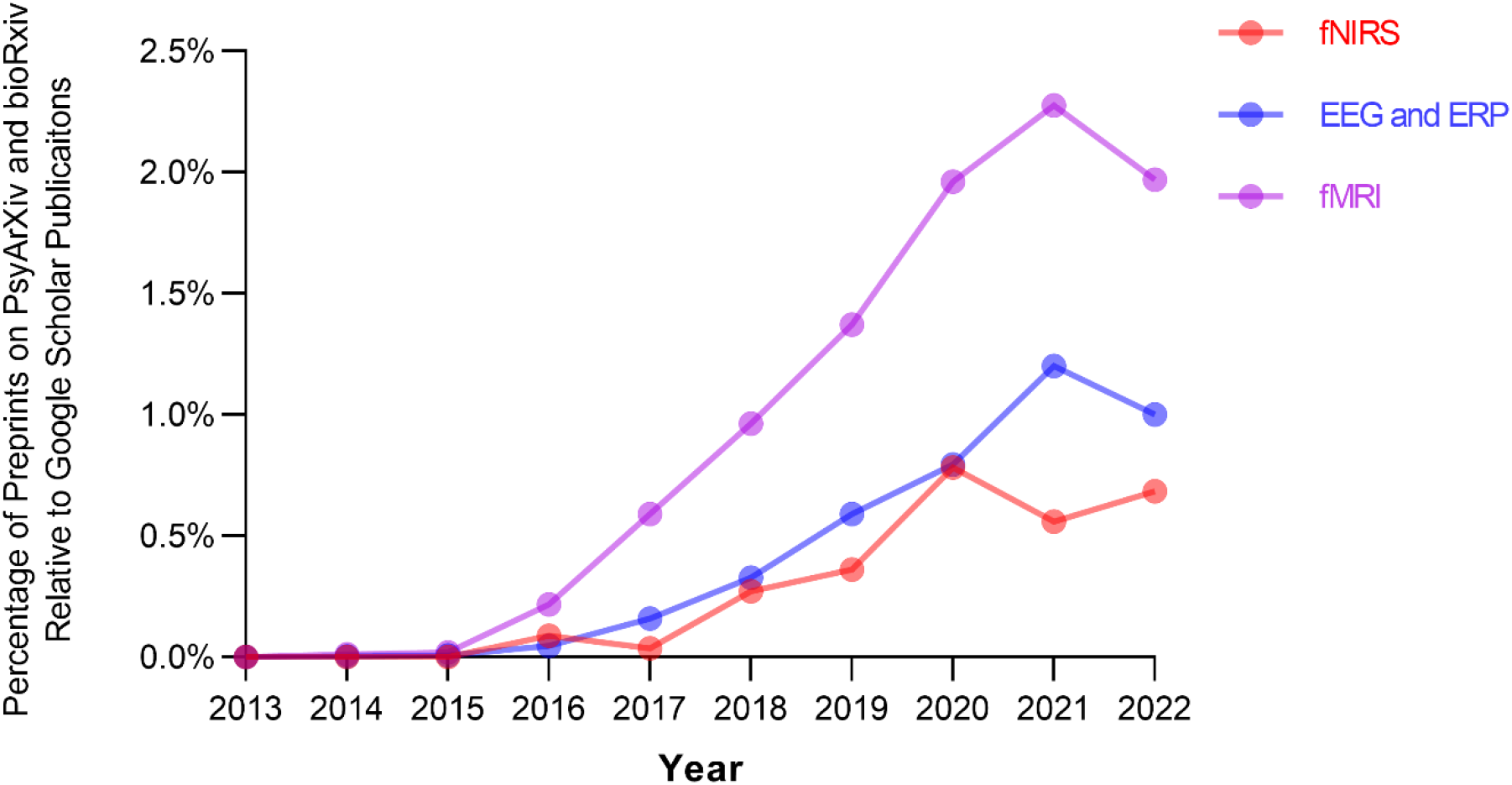
The ratio of preprints to publications for each imaging modality by year. *Note*. This search was conducted on 3/29/2022 using Share.osf.io (for PsyArXiv) and bioRxiv; please see [Supplementary Materials] for more details on these searches. The following search terms were used for PsyArXiv: fnirs OR “functional near infrared spectroscopy”, fmri OR “functional magnetic resonance imaging”, eeg OR erp OR electroencephalogram OR “event related potential”. The following search terms were used for bioRxiv: fnirs, fmri, eeg OR erp.

### Goals for the future

In the future, the field could consider creating a blinding system for preprints to keep the author’s identity anonymous while undergoing the peer review process and establishing more funding mechanisms to support publishing in open access journals.

## General conclusions and suggestions for the future

In this review, the important advances the fNIRS community has made in participating in the various initiatives of open science are highlighted. Although open science practices in the fNIRS field have become more common in recent years, there is ample room for wider adoption. To advance the fNIRS community’s participation in open science, the authors propose a number of general suggestions for the future. First, the authors propose the development of a working group for open science in fNIRS, in which researchers gather periodically to review the status of the field’s practices and compile recommendations and resources to help other researchers more easily participate in open science. Second, the authors recommend that open science education sessions are offered regularly, possibly at sfNIRS meetings and online webinars, to make researchers aware of best practices and provide resources to support the adoption of these practices. Third, the authors propose that open science practices should be specifically highlighted in job advertisements and valued in faculty searches to provide incentives for researchers to participate in open science. A related suggestion is that researchers may consider highlighting their open science practices on their CVs and on their lab websites by indicating how they have promoted open access by sharing code and stimuli (e.g., sharing links to their OSF or GitHub page) or have participated in transparent practices such as writing preregistrations and registered reports (e.g., putting an extra star next to publications where the methods and analyses were preregistered). Fourth, the authors propose contributing financial (e.g., writing into grant budget, soliciting donations) resources to support open science platforms (e.g., preregistration, preprint services, and data repositories). If these efforts are not supported, then there is the possibility the platforms will no longer be able to provide their specific services.

By adopting these suggestions, the fNIRS community can take steps towards advancing transparency, reproducibility, rigor, and efficiency in the field. Overall, we hope this review demystifies various stages of open science and clarifies the current state of open science in the fNIRS field.

## Supporting information

Registered Reports

## Acknowledgments

This work was supported by the National Institute of Mental Health (R01 MH078829; to CAN). This work was also supported by the National Institute of Child Health and Human Development (F32 HD105312-01A1; to CMK). The content is solely the responsibility of the authors and does not necessarily represent the official views of the National Institutes of Health.

## Footnotes

1 These are a few registries available: arXiv, Dryad, Figshare, GitHub, Mendeley, Neuroimaging Tools and Resources Collaboratory (NITRC), open science framework, rOpenSci.

2 These are a few data repositories: Data Archiving and Networked Services, Dataverse, fighshare, the NeuroImaging Tools & Resource Collaboratory, OpenNeuro, the Open Science Foundation, and Zenodo, among others

## References

APA Data Sharing Working Group. (2015). Data sharing: Principles and considerations for policy development. https://www.apa.org/science/leadership/bsa/data-sharing-report.pdf

Artemenko, C. (2021). Developmental fronto-parietal shift of brain activation during mental arithmetic across the lifespan: A registered report protocol. PLoS One, 16(8), e0256232. doi:10.1371/journal.pone.0256232

Blohowiak, B. B., Cohoon, J., de-Wit, L., Eich, E., Farach, F. J., Hasselman, F., … Lowrey, O. (2022). Badges to Acknowledge Open Practices. Retrieved from osf.io/tvyxz

Botvinik-Nezer, R., Holzmeister, F., Camerer, C. F., Dreber, A., Huber, J., Johannesson, M., … Adcock, R. A. (2020). Variability in the analysis of a single neuroimaging dataset by many teams. Nature, 582(7810), 84–88.

Bowring, A., Maumet, C., & Nichols, T. E. (2019). Exploring the impact of analysis software on task fMRI results. Human Brain Mapping, 40(11), 3362–3384. doi:https://doi.org/10.1002/hbm.24603

Chambers, C. D., & Tzavella, L. (2022). The past, present and future of Registered Reports. Nature Human Behaviour, 6(1), 29–42. doi:10.1038/s41562-021-01193-7

Collaboration, O. S. (2015). Estimating the reproducibility of psychological science. Science, 349(6251), aac4716. doi:doi:10.1126/science.aac4716

du Plessis, S., Oni, I. K., Lapointe, A. P., Campbell, C., Dunn, J. F., & Debert, C. T. (2022). Treatment of Persistent Postconcussion Syndrome With Repetitive Transcranial Magnetic Stimulation Using Functional Near-Infrared Spectroscopy as a Biomarker of Response: Protocol for a Randomized Controlled Clinical Trial. JMIR Res Protoc, 11(3), e31308. doi:10.2196/31308

Fanelli, D. (2012). Negative results are disappearing from most disciplines and countries. Scientometrics, 90(3), 891–904.

Flannery, J. (2020). fMRI Preregistration Template. Retrieved from osf.io/6juft

Gilmore, R. O., & Adolph, K. E. (2017). Video can make behavioral science more reproducible. Nature Human Behaviour, 1, 0128. https://doi.org/10.1038/s41562-017-0128

Gilmore, R. O., Diaz, M. T., Wyble, B. A., & Yarkoni, T. (2017). Progress toward openness, transparency, and reproducibility in cognitive neuroscience. Annals of the New York Academy of Sciences, 1396(1), 5–18. doi:https://doi.org/10.1111/nyas.13325

Govaart, G. H., Paul, M., & Schettino, A. (2021). Hack—Finalizing a preregistration template for ERP studies. Retrieved from osf.io/qt8vh

Grint, N. J., Johnson, C. B., Clutton, R. E., Whay, H. R., & Murrell, J. C. (2015). Spontaneous electroencephalographic changes in a castration model as an indicator of nociception: A comparison between donkeys and ponies. Equine Veterinary Journal, 47(1), 36–42. doi:https://doi.org/10.1111/evj.12250

Guérin, S. M. R., Vincent, M. A., Karageorghis, C. I., & Delevoye-Turrell, Y. N. (2021). Effects of Motor Tempo on Frontal Brain Activity: An fNIRS Study. Neuroimage, 230, 117597. doi:https://doi.org/10.1016/j.neuroimage.2020.117597

Gusenbauer, M., & Haddaway, N. R. (2020). Which academic search systems are suitable for systematic reviews or meta-analyses? Evaluating retrieval qualities of Google Scholar, PubMed, and 26 other resources. Research Synthesis Methods, 11(2), 181–217. doi:https://doi.org/10.1002/jrsm.1378

HEALthy Brain and Child Development Study (HBCD). (2022). Retrieved from https://heal.nih.gov/research/infants-and-children/healthy-brain#:~:text=Examples%20of%20HBCD%20Study%20activities,factors%20affect%20these%20developmental%20trajectories.

Henderson, E. L. (2022). A guide to preregistration and Registered Reports. MetaArXiv. doi:https://doi.org/10.31222/osf.io/x7aqr

Jiao, C., Li, K., & Fang, Z. (2022). Data sharing practices across knowledge domains: a dynamic examination of data availability statements in PLOS ONE publications. arXiv preprint arXiv:2203.10586.

Kaplan, R. M., & Irvin, V. L. (2015). Likelihood of null effects of large NHLBI clinical trials has increased over time. PLoS One, 10(8), e0132382.

Kidwell, M. C., Lazarević, L. B., Baranski, E., Hardwicke, T. E., Piechowski, S., Falkenberg, L.-S., … Nosek, B. A. (2016). Badges to Acknowledge Open Practices: A Simple, Low-Cost, Effective Method for Increasing Transparency. PLoS biology, 14(5), e1002456. doi:10.1371/journal.pbio.1002456

Li, X., Ai, L., Giavasis, S., Jin, H., Feczko, E., Xu, T., … Milham, M. P. (2021). Moving Beyond Processing and Analysis-Related Variation in Neuroscience. bioRxiv, 2021.2012.2001.470790. doi:10.1101/2021.12.01.470790

Mazziotti, R., Scaffei, E., Conti, E., Marchi, V., Rizzi, R., Cioni, G., … Baroncelli, L. (2022). The amplitude of fNIRS hemodynamic response in the visual cortex unmasks autistic traits in typically developing children. Translational psychiatry, 12(1), 53. doi:10.1038/s41398-022-01820-5

Meidenbauer, K. L., Choe, K. W., Cardenas-Iniguez, C., Huppert, T. J., & Berman, M. G. (2021). Load-dependent relationships between frontal fNIRS activity and performance: A data-driven PLS approach. Neuroimage, 230, 117795. doi:https://doi.org/10.1016/j.neuroimage.2021.117795

Munafò, M., & Neill, J. (2016). Null is beautiful: on the importance of publishing null results. In (Vol. 30, pp. 585–585): SAGE Publications Sage UK: London, England.

Nguyen, T., Hoehl, S., Bertenthal, B. I., & Abney, D. H. (2022). Coupling between prefrontal brain activity and respiratory sinus arrhythmia in infants and adults. Developmental cognitive neuroscience, 53, 101047. doi:https://doi.org/10.1016/j.dcn.2021.101047

Obels, P., Lakens, D., Coles, N. A., Gottfried, J., & Green, S. A. (2020). Analysis of Open Data and Computational Reproducibility in Registered Reports in Psychology. Advances in Methods and Practices in Psychological Science, 3(2), 229–237. doi:10.1177/2515245920918872

Petersen, I. T., Apfelbaum, K. S., & McMurray, B. (2022). Adapting open science and pre-registration to longitudinal research. Infant and Child Development, e2315.

Sarabipour, S., Debat, H. J., Emmott, E., Burgess, S. J., Schwessinger, B., & Hensel, Z. (2019). On the value of preprints: An early career researcher perspective. PLoS biology, 17(2), e3000151. doi:10.1371/journal.pbio.3000151

Simmons, J. P., Nelson, L. D., & Simonsohn, U. (2011). False-Positive Psychology:Undisclosed Flexibility in Data Collection and Analysis Allows Presenting Anything as Significant. Psychological Science, 22(11), 1359–1366. doi:10.1177/0956797611417632

Simon, D. A., Gordon, A. S., Steiger, L., & Gilmore, R. O. (2015). Databrary: Enabling sharing and reuse of research video. Proceedings of the 15th ACM/IEEE-CS Joint Conference on Digital Libraries, Knoxville, Tennessee, USA. https://doi.org/10.1145/2756406.2756951

Time to recognize authorship of open data. (2022). Nature, 604(7904), 8. https://doi.org/10.1038/d41586-022-00921-x

Wagenmakers, E.-J., Wetzels, R., Borsboom, D., van der Maas, H. L. J., & Kievit, R. A. (2012). An Agenda for Purely Confirmatory Research. Perspectives on Psychological Science, 7(6), 632–638. doi:10.1177/1745691612463078

Yücel, M., Lühmann, A., Scholkmann, F., Gervain, J., Dan, I., Ayaz, H., … Wolf, M. (2021). Best practices for fNIRS publications. Neurophotonics, 8(1), 012101. Retrieved from https://doi.org/10.1117/1.NPh.8.1.012101

