## Supplementary Materials for "Shedding Light on functional Near Infrared Spectroscopy and Open Science Practices"

### Supplementary Methods

#### Registered report search

The list of journals using the Registered Reports format was accessed via the Center for Open Science (COS) website on 3/20/2022 (https://www.cos.io/initiatives/registered-reports). For a paper to 'count' as publishing neuroimaging papers, it needed to involve the collection or analysis of EEG/ERP/fMRI/fNIRS. Specifically, secondary data analyses were counted, whereas meta-analyses/systematic reviews, mentions in theory papers, references to the modality in introduction or discussion sections, etc., were not counted. Additionally, instances of non-brain imaging (e.g., using fNIRS to assess blood flow in the arm) were not counted whereas papers with animal subjects were counted only if they were brain-based (e.g., Grint, Johnson, Clutton, Whay, & Murrell, 2015).

We used Google Scholar to determine if each of the journals listed on COS website met criteria for publishing an fNIRS or neuroimaging paper. To do this, we searched Google Scholar using these search terms: fnirs OR "functional near infrared spectroscopy" source:"journal". We then manually inspected the results (if any) to see if they met the criteria for counting. If no papers counted for fNIRS, we then checked for neuroimaging more broadly using the following search term: fnirs OR "functional near infrared spectroscopy" OR fmri OR "functional magnetic resonance imaging" OR eeg OR erp OR "electroencephalogram" OR "event related potential" source:"journal". Results of this search were again manually inspected. For a few journals with relatively generic titles (e.g. "Biolinguistics"), the search engine did not adequately filter results to the specific journal, and in these cases we searched the journal's website. Please see Online Supplemental Materials for more details on the journals for which this was the case, and for a representative publication (doi/url) for each journal that counted. It is also important to acknowledge the possible confounds of using the Google Scholar search engine (see Gusenbauer & Haddaway, 2020 for more details).

**Search terms**

When searching through GitHub repositories it was found that “ERP” meant to index “event related potential” was getting additional hits for referencing “enterprise resource planning”. We decided to keep ERP as a search term for all searchers as we believe there was no way to account for all other uses for the neuroimaging acronyms (e.g., fMRI, fNIRS, EEG, ERP).

#### Supplementary Tables

##### Supplementary Table 1

*Raw number of preregistrations for each imaging modality by year*

| Year | 2013 | 2014 | 2015 | 2016 | 2017 | 2018 | 2019 | 2020 | 2021 | 2022 | **Total** |
|---|---|---|---|---|---|---|---|---|---|---|---|
| fNIRS | 0 | 0 | 0 | 0 | 1 | 2 | 8 | 8 | 14 | 1 | **34** |
| fMRI | 1 | 1 | 14 | 8 | 27 | 35 | 54 | 129 | 131 | 36 | **436** |
| EEG | 0 | 4 | 4 | 10 | 19 | 45 | 67 | 142 | 168 | 45 | **504** |

*Note.* This search was conducted on 3/29/2022 using Share.osf.io (restricted to OSF Registries) and Google Scholar. This only includes preregistrations that have been made publicly available. The following search terms were used: fnirs OR “functional near infrared spectroscopy”, fmri OR "functional magnetic resonance imaging", eeg OR erp OR electroencephalogram OR "event related potential".

**Supplementary Table 2**

*The number of codes available on GitHub for each imaging modality separated by coding language.*

| Programming Language | fNIRS | EEG/ERP | fMRI |
|---|---|---|---|
| Matlab | 92 | 1956 | 910 |
| Python | 49 | 5318 | 804 |
| Jupyter Notebook | 24 | 1817 | 473 |
| Shell | 0 | 0 | 188 |
| R | 9 | 0 | 157 |
| Java | 7 | 3810 | 0 |
| PHP | 0 | 2363 | 0 |
| C | 5 | 0 | 15 |
| HTML | 4 | 2174 | 103 |
| JavaScript | 4 | 4207 | 25 |
| Smarty | 3 | 0 | 0 |
| C# | 2 | 1805 | 0 |
| C++ | 0 | 0 | 29 |
| CSS | 0 | 894 | 0 |
| TeX | 0 | 0 | 18 |
| TypeScript | 0 | 753 | 0 |
| **Total** | **277** | **39808** | **3419** |

*Note.* This search was conducted on 3/29/2022 using GitHub. The following search terms were used: fnirs OR "functional near infrared spectroscopy", fmri OR "functional magnetic resonance imaging", eeg OR erp OR electroencephalogram OR "event related potential"

**Supplementary Table 3**

*Raw number of preprints for each imaging modality by year for PsyArXiv and bioRxiv.*

| Year | 2013 | 2014 | 2015 | 2016 | 2017 | 2018 | 2019 | 2020 | 2021 | 2022 | **Total** |
|---|---|---|---|---|---|---|---|---|---|---|---|
| ***PsyArXiv, bioRxiv (total)*** | | | | | | | | | | | |
| fNIRS | 0,0<br>(0) | 0,0<br>(0) | 0,0<br>(0) | 0,2<br>(2) | 0,1<br>(1) | 4,5<br>(9) | 6,8<br>(14) | 9,27<br>(36) | 8,24<br>(32) | 3,5<br>(8) | **30,72**<br>**(102)** |
| fMRI | 0,1<br>(1) | 0,6<br>(6) | 0,11<br>(11) | 1,109<br>(110) | 20,282<br>(302) | 54,429<br>(483) | 75,560<br>(635) | 99,682<br>(781) | 93,726<br>(819) | 29,172<br>(201) | **371,2978**<br>**(3349)** |
| EEG | 0,0<br>(0) | 0,3<br>(3) | 0,9<br>(9) | 0,56<br>(56) | 21,166<br>(187) | 45,286<br>(331) | 85,470<br>(555) | 117,515<br>(632) | 126,557<br>(683) | 39,163<br>(202) | **433,2225**<br>**(2658)** |

*Note.* This search was conducted on 3/29/2022 using Share.osf.io (for PsyArXiv) and bioRxiv; The following search terms were used for PsyArXiv: fnirs OR “functional near infrared spectroscopy”, fmri OR "functional magnetic resonance imaging", eeg OR erp OR electroencephalogram OR "event related potential". The following search terms were used for bioRxiv: fnirs, fmri, eeg OR erp. We planned to use Share.osf.io for both of these servers, but while compiling numbers we discovered that SHARE does not list preprints added to bioRxiv Feb. 2019-present; further, bioRxiv does not allow searches to include both “stand-alone phrases” and Boolean operations, so we chose to use the individual abbreviations and restricted the results to hits in the “Abstract or Title”.
